# Local RhoA Activation Induces Cytokinetic Furrows Independent of Spindle Position and Cell Cycle Stage

**DOI:** 10.1101/043836

**Authors:** Elizabeth Wagner, Michael Glotzer

## Abstract

Cytokinetic cleavage furrows assemble during anaphase at a site that is dictated by the position of the spindle. The GTPase RhoA promotes contractile ring assembly and furrow ingression during cytokinesis. While many factors that regulate RhoA during cytokinesis have been characterized, the spatiotemporal regulatory logic remains undefined. It is not known whether a local zone of RhoA activity is sufficient to induce furrow formation or whether the spindle modulates furrow assembly through other pathways. Similarly, it is not known whether the entire cortex is responsive to RhoA, nor whether contractile ring assembly is cell cycle regulated. Here, we have developed an optogenetic probe to gain tight spatial and temporal control of RhoA activity in mammalian cells and demonstrate that cytokinetic furrowing is primarily regulated at the level of RhoA activation. Light-mediated recruitment of a RhoGEF domain to the plasma membrane leads to rapid activation of RhoA, leading to assembly of cytokinetic furrows that partially ingress. Furthermore, furrow formation in response to RhoA activation is not spatially or temporally restricted. RhoA activation is sufficient to generate furrows at both the cell equator and at cell poles, in both metaphase and anaphase. Remarkably, furrow formation can be initiated in rounded interphase cells, but not adherent cells. These results indicate RhoA activation is sufficient to induce assembly of functional contractile rings and that cell rounding facilitates furrow formation.

## Introduction

Cytokinesis is the final stage of cell division in which an actomyosin-based contractile ring physically divides the cell into two genetically equivalent daughter cells. Our understanding of cytokinesis has been greatly influenced by classical experiments in which spindles and/or blastomeres were repositioned or micro-manipulated. These perturbations demonstrated that the spindle induces furrow formation during a specific time interval following anaphase onset (Rappaport, 1985). At a molecular level, the small GTPase, RhoA, serves as an essential and dosage sensitive regulator of cleavage furrow formation in metazoan cells (Fededa and Gerlich, 2012; Loria et al., 2012; Kishi et al., 1993). RhoA behaves like a molecular switch that is active when bound to GTP. When active, RhoA binds to downstream effectors including the formin, mDia2, to induce F-actin assembly (Watanabe et al., 2008; Otomo et al., 2005) and Rho Kinase, ROCK, to activate non-muscle myosin II (Kosako et al., 2000). Through these and other effectors, RhoA regulates the dynamic changes in actomyosin required for contractile ring assembly and cleavage furrow formation.

RhoA activation during cytokinesis is spatially and temporally regulated and dependent upon the RhoGEF Ect2 (Tatsumoto et al., 1999). Ect2 localization and activation are regulated by a phospho-dependent interactions with centralspindlin, a protein complex that accumulates on the spindle midzone during anaphase (Yüce et al., 2005; Wolfe et al., 2009; Petronczki et al., 2007; Burkard et al., 2009; Zhang and Glotzer, 2015)(Supplemental Fig. 1). This complex also accumulates in small amounts on the cortex where it directs local accumulation of RhoA activation (Basant et al., 2015). Despite extensive research, several major questions concerning the regulation of cytokinesis remain unanswered. Is local activation of RhoA is sufficient to generate a cleavage furrow or are other factors required in parallel to RhoA for furrow formation? Are there spatial or temporal requirements for RhoA-mediated contractile ring assembly and furrow formation?

Answers to these fundamental questions require the ability to spatially and temporally manipulate cytokinesis at the molecular level, in particular, at the level of RhoA activation. Optogenetic tools afford the possibility of precise control of protein localization and, in many cases, control of localization allows control of protein activity (Strickland et al., 2012; Toettcher et al., 2013). We have engineered an optogenetic tool that affords local control of RhoA activity and have used this to demonstrate that local activation of RhoA is sufficient to direct cleavage furrow formation.

## Results and Discussion

### Light-mediated control of RhoA activity

Previous iterations of the two component optogenetic system TULIPs utilized a membrane-targeted photosensitive domain, LOVpep, in conjunction with a second bioengineered tag, ePDZ-b1, that binds to LOVpep in a light-dependent manner (Strickland et al., 2012). Here, we substituted the ePDZ-b1 tag with a tandem PDZ tag which is functional in more diverse protein fusions. To manipulate RhoA activation with light, we fused the tandem PDZ tag to the highly specific RhoA guanine nucleotide exchange factor (GEF) LARG (Jaiswal et al., 2011), to create a construct we refer to as optoGEF (Fig. 1A). To reduce basal activity, only the catalytic GEF DH domain was included. GFP-tagged LOVpep was localized to the plasma membrane by fusing it to the transmembrane receptor Stargazin (Stargazin-GFP-LOVpep). A digital micro mirror device (DMD) was used to illuminate arbitrarily defined regions of the cell with 405 nm light. Illumination of adherent cells expressing these constructs resulted in light-mediated local recruitment of optoGEF (Fig. 1B-C and Supplementary Video 1). Recruitment also leads to local accumulation of myosin (mCherry-MLC) and F-actin (mApple-actin)(Fig. 1C-G and Supplementary Video 2 and 3) within 20-40 secs. When illumination ceases, the local increase in GEF recruitment is rapidly lost, consistent with the thermal reversion of LOVpep into the dark state (Strickland et al., 2012) (t_½_ = 80 sec)(Fig. 1C). The local increases in actin and myosin are lost with roughly similar kinetics (Fig. 1E,G), suggesting that RhoA activation is not self-sustaining and suggest the existence of RhoGAPs that rapidly inactivate the ectopically activated RhoA.

**Figure 1:**
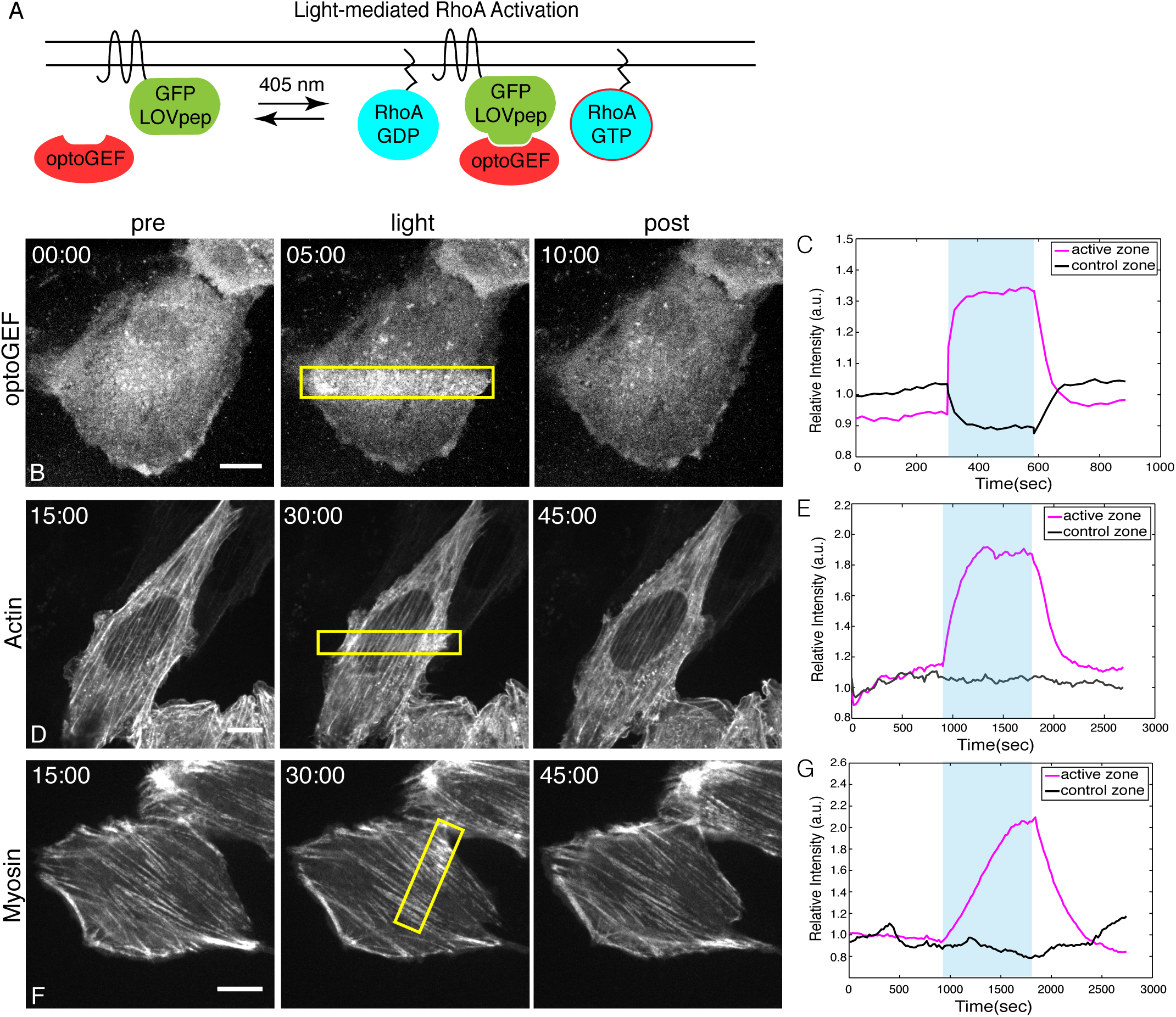
Light-mediated activation of RhoA. (A) TULIPs-mediated activation of RhoA by light-directed recruitment of optoGEF. Local illumination of NIH3T3 cells with 405 nm light (yellow boxes) induces recruitment local recruitment of optoGEF (B), F-actin polymerization (D), and myosin accumulation (F). Quantification of the relative increase in intensity in the region of activation (magenta) vs. a control region (black) for optoGEF (C), mApple-actin (E), and mCherry-MLC (G) over time. During the photoactivation period, designated by the blue box, cells were locally illuminated with 960 ms pulse every 20 sec and 561 images were taken every 20 secs for the entire experiment. All scale bars are 10 *μ*m.

### Local activation of RhoA is sufficient to initiate furrow formation at the midzone of anaphase cells

A central question we sought to answer was whether a local zone of RhoA activation is sufficient to form a cleavage furrow. The least stringent initial test of this model is to determine whether, in a cell progressing normally through anaphase, light-induced RhoA activation can substitute at the equator for the endogenous pathway. The mitotic kinase, Polo-like kinase 1 (Plk1), is required for the phospho-dependent interaction between centralspindlin and Ect2; Plk1 inhibition therefore precludes RhoA activation and furrow formation (Wolfe et al., 2009; Petronczki et al., 2007). This provides an appropriate context for examining whether light-mediated activation of RhoA could be sufficient to induce furrow formation at the midzone.

To generate non-contractile anaphase cells, HeLa cells were arrested in metaphase using a low dose of nocodazole (30 ng/ml) for 4-5 hrs. Release of the block allows cells to initiate mitotic exit. Expression of optoGEF components does not impair cytokinesis (Fig. 2B). Plk1 inhibitor (BI 2536, 200 nM) was added 30 min after release to block endogenous RhoA activation while permitting anaphase onset (Fig. 2A) (Petronczki et al., 2007). In the absence of optoGEF recruitment, cells remain non-contractile and furrows do not form (Fig. 2C). Photoactivation of an equatorial zone induced recruitment of optoGEF to a finely limited region. Light-mediated activation of RhoA at the midzone was sufficient to initiate cleavage furrow formation (Fig. 2D and Supplementary Video 4). Furrows resulting from optoGEF recruitment require the RhoA pathway as furrowing was inhibited by Rho kinase inhibitor (10 *μ*M Y27632) (Supplemental Fig. 2) and recruitment of PDZ_2_-mCherry failed to induce furrow formation (Fig. 2E). The initial rate of ingression was similar to that of cells dividing normally in the absence of BI 2536 (Fig. 2F). However, light-induced furrows do not fully ingress (~34%, n=32). In the absence of continued photoactivation, these furrows regress, indicating that RhoA activation during cytokinesis is not self-sustaining, at least in the presence of Plk1 inhibitor. In addition to regulating the phospho-dependent Ect2:centralspindlin interaction, Plk1 may have additional roles in promoting furrow ingression (Wolfe et al., 2009; Niiya et al., 2006; Lowery et al., 2007; Neef et al., 2003; Liu et al., 2004). To generate non-contractile anaphase cells without inhibiting Plk1, cells were depleted of HsCyk4. Light-mediated activation of RhoA in Cyk4-depleted cells resulted in similar ingression as in Plk1-inhibited cells (Supplemental Fig. 3), suggesting that Plk1 functions primarily upstream of RhoA activation.

**Figure 2:**
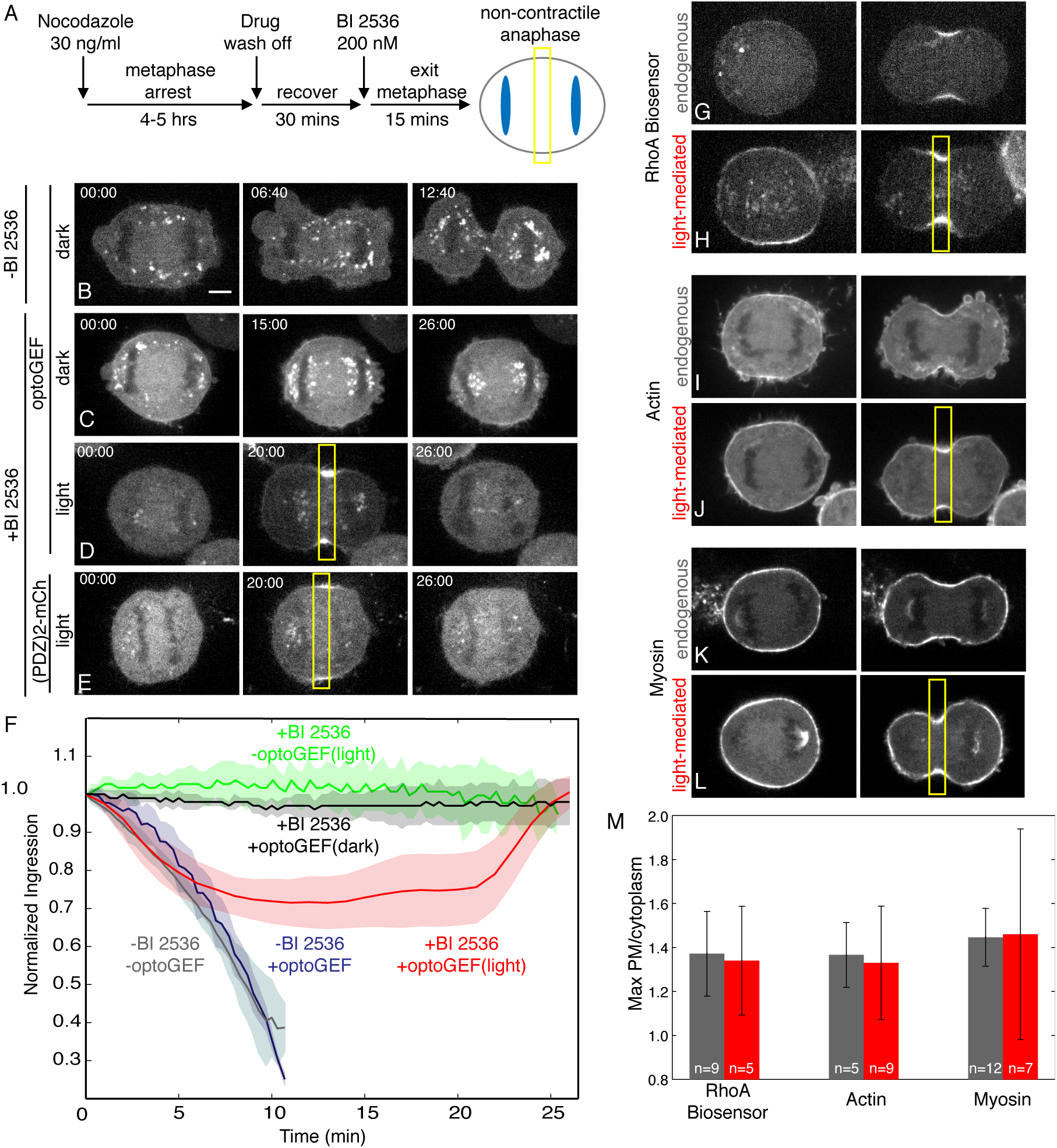
Local activation of RhoA is sufficient to induce furrow formation in anaphase cells at the midzone. (A) Experimental design to generate non-contractile anaphase HeLa cells. (B) Images of control HeLa cell expressing optoGEF dividing normally with no illumination (dark). Images of non-contractile anaphase HeLa cells (200 nM BI 2536) expressing optoGEF (C,D) or PDZ2-mCh (E) either with (D,E) or without local illumination (C). (F) Average normalized furrow ingression overtime of HeLa cells dividing normally (_-_BI 2536) with (blue, n=7) and without (grey, n=10) optoGEF expression, non-contractile anaphase cells (+BI 2536) expressing optoGEF with (red, n=32) and without local illumination (black, n=6), and local illuminated non-contractile anaphase cells (+BI 2536) expressing PDZ2-mCherry (green, n=5). Images of RhoA biosensor (AHDPH-mCherry) (G,H), F-actin (mApple-actin)(I,J), and myosin (mApple-MLC) (K,L) levels in cells dividing normally and during light-mediated furrow formation. (M) Maximum PM/cytoplasm ratio of RhoA activation, Actin, and Myosin levels in endogenous (grey) vs. light-mediated (red) furrows. For all photoactivation experiments, cells were locally illuminated at the midzone (yellow box) every 20 secs with a 960ms pulse for 20 mins followed by 10 mins without local activation and 561 nm images were taken every 20 secs throughout the experiment. Scale bar is 5 *μ*m.

To test whether incomplete ingression might be a consequence of insufficient levels of RhoA activation, we used a RhoA biosensor (Piekny and Glotzer, 2008) to compare the levels of active RhoA resulting from light-mediated GEF recruitment to that of normally dividing cells. The distribution and level of active RhoA are comparable to those in normally dividing cells (Fig. 2G,H,M). In addition, two downstream effectors of RhoA, F-actin and myosin, accumulate at levels comparable to those observed in normally dividing cells (Fig. 2I-M). Therefore incomplete ingression does not appear to caused by dramatically reduced activation of RhoA or its key effectors.

### The anaphase cortex is uniformly responsive to RhoA activation

To more stringently test whether local RhoA activation is sufficient to induce furrow formation, we exploited the flexibility that optogenetic tools provide to determine whether the response to RhoA activation is spatially modulated in the anaphase cortex. The anaphase spindle has been shown to play both positive and negative roles in directing RhoA activation and cleavage furrow formation. The central spindle promotes local RhoA activation and cleavage furrow formation at the midzone. Conversely, dynamic astral microtubules play a role in inhibiting cortical contractility in the polar regions (Werner et al., 2007; Zanin et al., 2013). Furthermore, the spindle may provide additional positive cues in addition to locally regulating RhoA, such as directed membrane trafficking (Drechsel et al., 1997) and local accumulation of mitotic kinases (Golsteyn et al., 1995; Adams et al., 2001).

Using the same experimental design where endogenous RhoA activation pathway is inhibited by the Plk1 inhibitor, we induced local RhoA activation in a zone encompassing the cell poles. RhoA activation is sufficient to initiate furrow formation at this site. These furrows behave similarly to those at induced at the midzone, ingressing ~36% at a constriction rate of 4.5 *μ*m/min (Fig. 3A-C and Supplementary Video 5). To directly compare the response to RhoA activation in the midzone vs. poles we simultaneously activated both regions in the same cell. Concurrent illumination of the midzone and poles induces furrows in both regions that ingress at similar rates and to similar extents (Fig. 3D-F and Supplementary Video 6). Due to the absence of midzone derived cues at poles, these results demonstrate that RhoA activation alone is sufficient to induce cleavage furrow ingression and the entire cortex is equally responsive to RhoA activation. This also suggests that astral inhibition does not principally act at the level of active RhoA or its downstream effectors. However, one caveat is that Plk1 could suppress astral-mediated inhibition, but we consider this to be unlikely, particularly because Plk1 normally concentrates at the midzone (Golsteyn et al., 1995).

**Figure 3:**
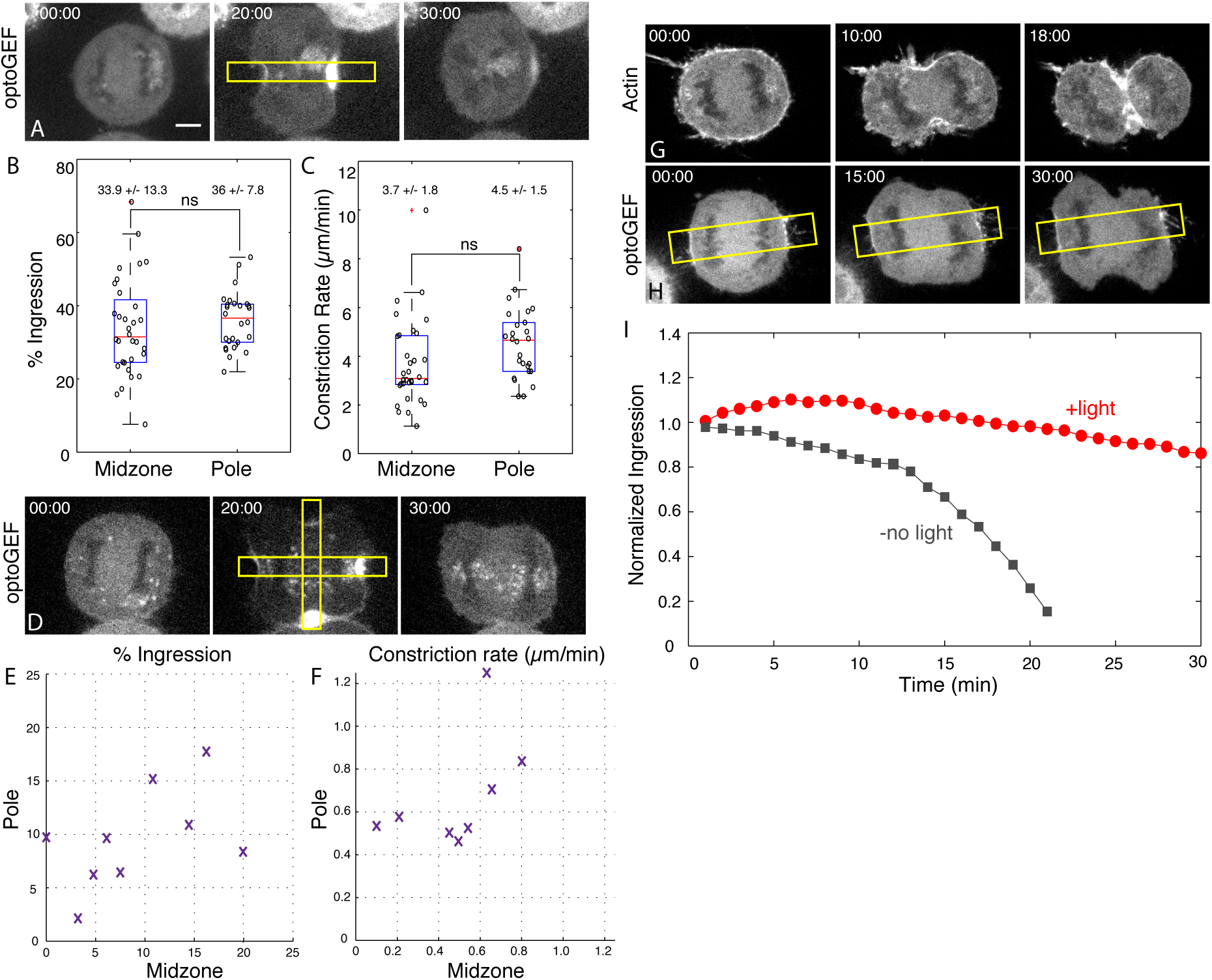
The anaphase cortex is uniformly responsive to RhoA activation. (A) Images of non-contractile anaphase HeLa cells (200 nm BI 2536) expressing optoGEF locally activated (yellow boxes) at the polar region. Quantification of the percent ingression (B) and Constriction Rate (*μ*m/min) (C) of light-mediated furrows formed in midzone vs. pole. (D) Images of non-contractile anaphase HeLa cells (200 nm BI 2536) expressing optoGEF simultaneously activated (yellow boxes) at the midzone and polar regions (yellow boxes). Quantification of the % Ingression (E) and Constriction Rate (*μ*m/min) (F) measured at the midzone vs pole for individual cells (n=8). (G) Images of HeLa cell expressing mApple Actin undergoing cytokinesis (-BI 2536). (H) Images of a HeLa cell expressing optoGEF undergoing cytokinesis (-BI 2536) with polar photoactivation. (I) Ingression kinetics of the endogenous furrow with (red) and without (grey) polar photoactivation. For all photoactivation experiments, cells were locally illuminated at the designated regions (yellow boxes) every 20 secs (960 ms pulse) for 20 mins or 30 mins (H) followed by 10 mins without local activation and 561 nm images were taken every 20 secs throughout the experiment. Scale bar is 5 *μ*m.

As optoGEF can induce furrowing at the poles, we also assessed the ability of exogenous RhoA activation to compete with the endogenous pathway. Polar recruitment of optoGEF dramatically slows ingression of the endogenous furrow; in some cases blocking furrow initiation altogether (Fig. 3G-I and Supplementary Video 7). Although we do not observe deep furrows in the poles in these perturbations, we observe local flattening and suppression of blebbing in the zone of activation, indicating polar activation of RhoA. As RhoA activation occurs despite Plk1 being uninhibited, this further suggests that astral inhibition is not regulated by Plk1.

### The response to RhoA activation is not strongly regulated during cell cycle

We next sought to determine if the response to RhoA activation is temporally regulated. Anaphase onset results in a dramatic increase of cortical contractility. Because the upstream RhoA activation pathway is largely suppressed by Cdk1 activity in metaphase (Yüce et al., 2005), it is not known whether active RhoA or its downstream effectors are subject to cell cycle regulation. Cells were arrested using a low dose of nocodazole (30 ng/ml) for 4-5 hrs and released for 20 mins to allow assembly of astral microtubules. Prior to mitotic exit, RhoA was locally activated and furrow formation was induced. The response to local RhoA activation was similar to that induced during anaphase. Furrows ingressed ~33% at a rate of ~4.8 *μ*m/min (Fig. 4A,B, Supplemental Fig. 5, and Supplementary Video 8). Thus, there does not appear to be potent metaphase-specific regulation of active RhoA or its downstream effectors. OptoGEF-induced furrow formation occurs irrespective of whether the zone of illumination was parallel or perpendicular to the metaphase plate, indicating that the response to RhoA activation during metaphase is not spatially regulated.

**Figure 4:**
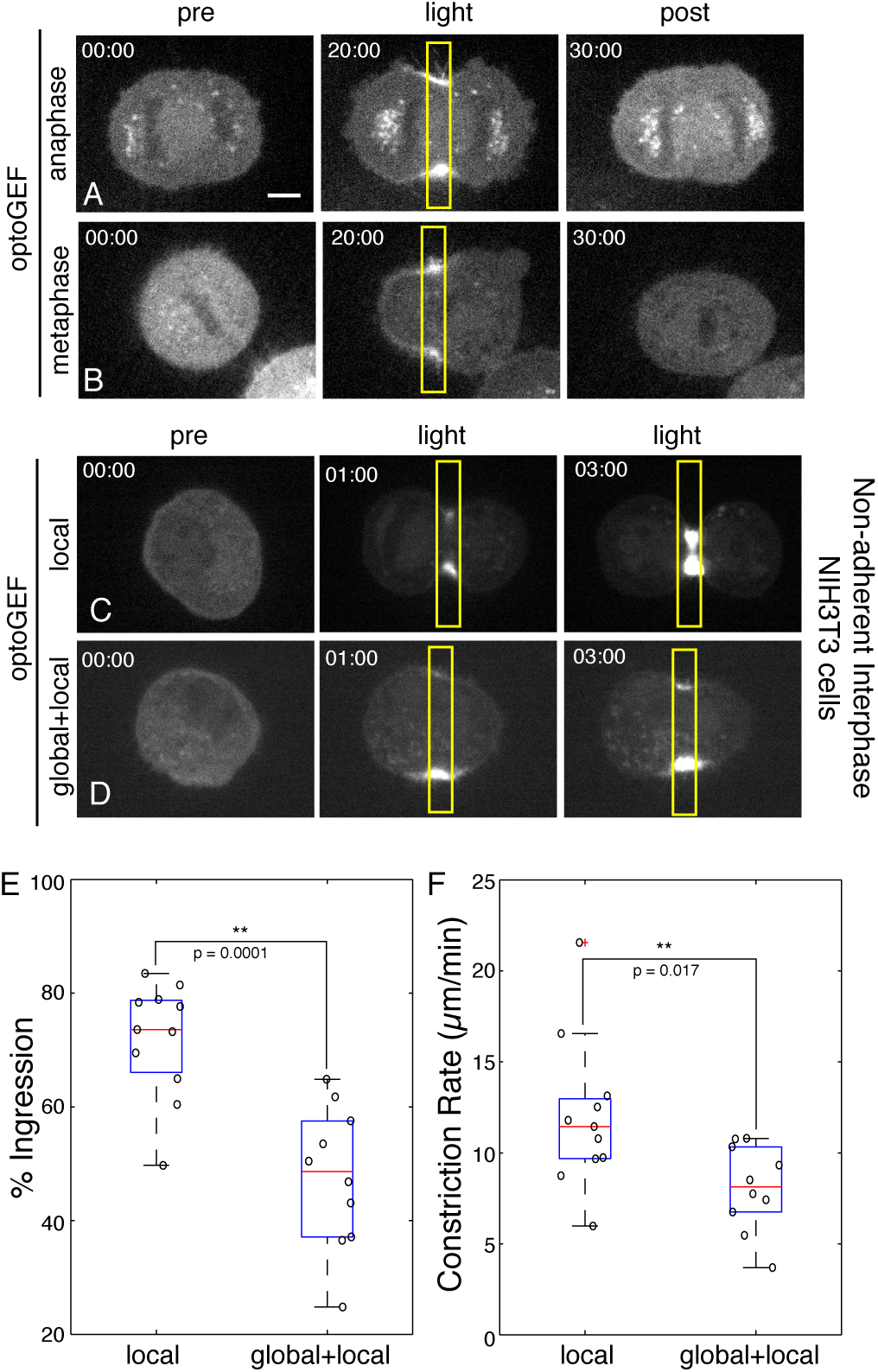
The response to RhoA activation is not strongly cell cycle regulated. Images of non-contractile anaphase HeLa cell (200 nM BI 2536) (A) and untreated metaphase cells (B) locally illuminated (yellow box) every 20 secs (960 ms pulse) for 20 mins followed by 10 min without photoactivation and 561 nm images were taken every 20 secs. Non-adherent interphase NIH3T3 cells locally illuminated (960 ms pulse) (C) or in combination with short global illumination (10 ms pulse) (D) every 20 secs for 20 mins. Quantification of % Ingression (E) and Constriction Rate (*μ*m/min) (F) of non-adherent interphase NIH3T3 cells illuminated with a local or local+ global pulse. Scale bar is 5 *μ*m.

The consistent response to RhoA activation irrespective of mitotic stage or spindle position prompted us to investigate whether local activation of RhoA can induce furrow formation during interphase. As shown in Figure 1, local activation of RhoA in adherent interphase cells generates F-actin and myosin II accumulation but does not induce furrowing. However, cells entering the mitotic cycle frequently remodel adherence to the substrate and to neighboring cells, allowing them to round up (Meyer et al., 2011; Stewart et al., 2011). We hypothesized that a decrease in cell adhesion would alter the response to local RhoA activation in interphase cells. To induce cell detachment and rounding we treated NIH3T3 cells with trypsin-EDTA prior to replating. Upon local illumination, cells rapidly form furrows which ingresses to 72% of completion (Fig. 4C and Supplementary Video 9). Similar results were observed in nonadherent interphase HeLa cells (Supplemental Fig. 5). These results strongly demonstrate that local activation of RhoA is sufficient to self assemble a contractile ring and no mitotic specific factors are required for cleavage furrow formation.

### Cortical tension modulates furrow ingression in response to local RhoA activation

Exogenous activation of RhoA is sufficient to initiate furrow formation throughout the cell cycle, however, furrows ingress further in non-adherent interphase cells compared to those induced in metaphase or anaphase. The distinct mechanical properties of mitotic vs rounded interphase cells may explain this differential response. Mitotic entry induces an isotropic increase cortical tension and hydrostatic pressure concomitant with cell rounding; the rounding pressure is approximately 3-fold higher in metaphase as compared to non-adherent interphase cells (Stewart et al., 2011). The enhanced furrow ingression in non-adherent interphase cells could therefore reflect their increased compliance relative to mitotic cells.

We therefore modulated the levels of global cortical tension and assessed whether this altered the response to local RhoA activation in non-adherent interphase cells. To increase the levels of global cortical tension we globally activated RhoA. By varying the duration of the illumination pulse, the levels of GEF recruitment can be finely tuned (Supplemental Fig. 6). Therefore, in addition to the standard local photoactivation pulse (960 ms) we also globally illuminated cells with a short pulse (10 ms) to modestly increase global cortical tension. In comparison to local activation alone, global activation induces a 1.5-fold decrease in both the extent of ingression and the constriction rate (Fig. 4D-F and Supplementary Video 10), demonstrating that cortical compliance impacts furrow ingression.

Extensive genetic analysis demonstrates that RhoA activation is necessary for contractile ring assembly and cleavage furrow formation during cytokinesis (Fededa and Gerlich, 2012). However, whether RhoA activation is the primary control point for cleavage furrow assembly has not been previously examined. An optogenetic approach to control RhoA activation with high spatial and temporal precision has allowed us to molecularly dissect the regulatory logic underlying furrow formation. Exogenous activation of RhoA is sufficient to rescue furrow formation at the midzone and the anaphase cortex is uniformly responsive to RhoA activation. These results demonstrate that neither the spindle midzone nor astral microtubules directly regulate active RhoA or its downstream effectors during furrow initiation. RhoA activation is also sufficient to induce contractile ring assembly in non-adherent interphase cells, indicating that no mitosis-specific factors are required for cleavage furrow formation. Decreased cell adhesion and rounding are commonly observed in dividing cells both in culture and within tissues (Meyer et al., 2011). As active RhoA does not induce furrows in adherent cells, our results suggest that cell detachment and rounding may be important for the ability of active RhoA to induce cleavage furrow formation.

The response to RhoA activation is not strongly cell cycle regulated, as we observe equal furrow formation in metaphase and anaphase cells. The enhanced furrow ingression observed in non-adherent interphase cells may be due to their reduced cortical tension as compared to mitotic cells (Stewart et al., 2011). Reduced levels of active RhoA or its effectors do not appear to be responsible for the incomplete ingression of furrows in anaphase cells. We speculate that the central spindle provides a pool of Ect2-centralspindlin complex that supports complete furrow ingression by activating RhoA with high spatial precision. Accumulating evidence indicates centralspindlin, Ect2, and RhoA are engaged in a positive feedback loop that is predicted to strengthen and sharpen the zone of RhoA activation (Bement et al., 2015; Zhang and Glotzer, 2015). We speculate that this, and perhaps other, feedback mechanisms drive complete furrow ingression in anaphase cells.

## Materials and Methods

### Cell culture and Drug Treatment

HeLa and NIH3T3 (ATCC) cells were grown in Dulbecco’s modified Eagles medium (Sigma) supplemented with 10% fetal bovine serum (Hyclone; Thermo Fisher Scientific), 0.2 mM L-glutamine (Invitrogen), and 1% penicillin/streptomycin (Invitrogen). Where indicated, endogenous HsCyk4 was depleted as described previously (Yüce et al., 2005). For HeLa cell experiments, cells were plated onto glass coverslips one day before transfection with plasmid DNA and siRNAs (where applicable) using Lipofectamine 2000 (Invitrogen). For all mitotic cell experiments, 24 h post transfection, cells were arrested in prometaphase with a 30 ng/ml Nocodazole (Sigma) for 4-6 hours. The nocodazole was then washed off and cells were incubated for 30 mins to allow them to recover and exit metaphase. BI-2536 (200 nM, generously provided by Norbert Kraut, Boehringer Ingelheim) was then added to block RhoA activation as cells entered anaphase. For NIH3T3 cell experiments, plasmids were transiently transfected using a Neon electroporation system (Invitrogen). For nonadherent cell experiments, cells were trypsinized using 0.05% trypsin-EDTA (Sigma), washed, replated onto glass coverslips and incubated for 5-10 mins to allow minimal adhesion prior to imaging.

### Constructs

The optogenetic membrane tether consisted of Stargazin-GFP-LOVpep fusion. Full-length Stargazin (a gift from A Karginov, University of Illinois Chicago, Chicago IL) was used. The LOVpep variant used is T406A, T407A, I532A (Strickland et al., 2012). The optoGEF construct consists of a (PDZ)_2_-mCherry-LARG(DH) fusion protein. The PDZ domain is derived from Erbin protein (Skelton et al., 2003) (a gift from S. Koide, University of Chicago, Chicago IL), which was fused to the DH domain (aa 766-997) of the RhoGEF LARG (NM_015313.2). Flexible linkers (SAGG_3_ and SAGG_5_, respectively) were placed between the PDZ domains and between mCherry and the LARG DH domain. OptoGEF^YFP^ was constructed in an identical manner to optoGEF with YFP replacing mCherry. This construct was used in experiments where the effects on various downstream markers were visualized. To examine the levels of RhoA activation we used the RhoA biosensor (AHDPH-mCherry) (Piekny and Glotzer, 2008). To examine effects on the actin and myosin networks, we used mApple-Actin and mApple-MLC constructs (gifts from M Davidson, U. Florida, Gainesville FL).

### Live Cell Imaging and Photoactivation

Glass coverslips were placed into an imaging chamber with media supplemented with 10 mM HEPES and 3% Oxyrase (Oxyrase Inc., Mansfield, OH) and maintained at 37°C. Cells were imaged using a 60X 1.49 NA ApoTIRF oil-immersion objective on a Nikon Ti-E inverted microscope (Nikon, Melville, NY). The microscope is fitted with a spinning disk confocal (CSU-X1; Yokogawa Electric, Musashino, Tokyo, Japan), illuminated with a laser merge module containing 491 nm and 561 nm lasers (Spectral Applied Research, Richmond Hill, Ontario, Canada). Images were acquired with Coolsnap HQ2 CCD camera (Photometics). A Mosaic digital micromirror-based device (DMD) equipped with a 100 mW 405 nm laser (Photonic Instruments) was used for photoactivation. MetaMorph acquisition software (Molecular Devices, Eugene, OR) was used to control the microscope hardware. The photoactivation laser was electronically attenuated and optically filtered such that the total incident light if the entire DMD was illuminated was < 1 *μ*J/s. The defined ROI was illuminated for 960 ms every 20 secs for the defined photoactivation period. When the entire cell was globally activated, an additional 10 ms exposure was added. To select cells for study, confocal images were acquired with 491 nm light followed by 561 nm light and robust recruitment of optoGEF was visually confirmed. To examine effects on RhoA activation levels (AHD-mCherry), actin (mApple-actin), or myosin (mApple-MLC), these constructs were co-expressed with Stargazin-GFP-LOVpep and optoGEF^YFP^. Cells strongly expressing both the reporter and the membrane tether were chosen for study.

## Acknowledgements

The authors thank Patrick Oakes and Margaret Gardel for generous access to their imaging system and for sharing routines for quantification, Andrei Karginov, Shohei Koide, and Mike Davidson for generous gifts of plasmids. This work was supported by NIGMS GM085087 to MG and EKW was supported by NIGMS T32 GM007183. We also thank our colleagues in MGCB for advice and support.

## Supplemental Figures

**Supplemental Figure 1:**
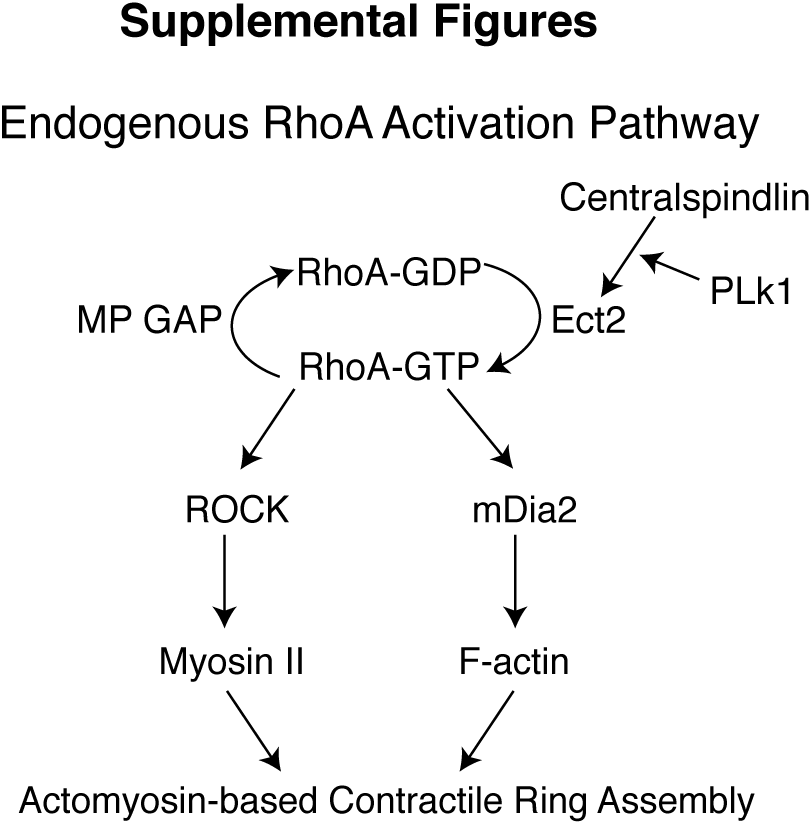
Endogenous pathway for RhoA activation during cytokinesis. (A) Schematic diagram depicting the pathway that promotes RhoA activation at the equatorial cortex during anaphase.

**Supplemental Figure 2:**
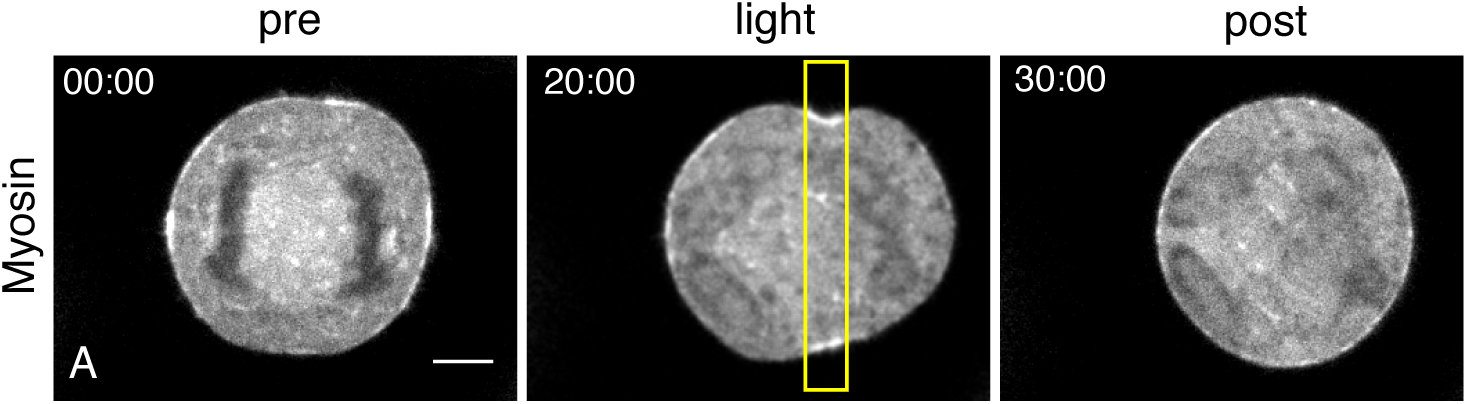
ROCK inhibition blocks light-mediated furrow formation. (A) Following release from nocodazole, HeLa cells were incubated in 10 μM ROCK inhibitor Y27632 for 30 mins. 200 nM BI 2536 was then added to generate non-contractile anaphase cells. Cells were locally illuminated (yellow box) every 20 secs (960 ms pulse) for 20 mins followed by 10 mins without local illumination. Little to no furrow ingression was observed in 4/4 cells. Scale bar is 5 *μ*m.

**Supplemental Figure 3:**
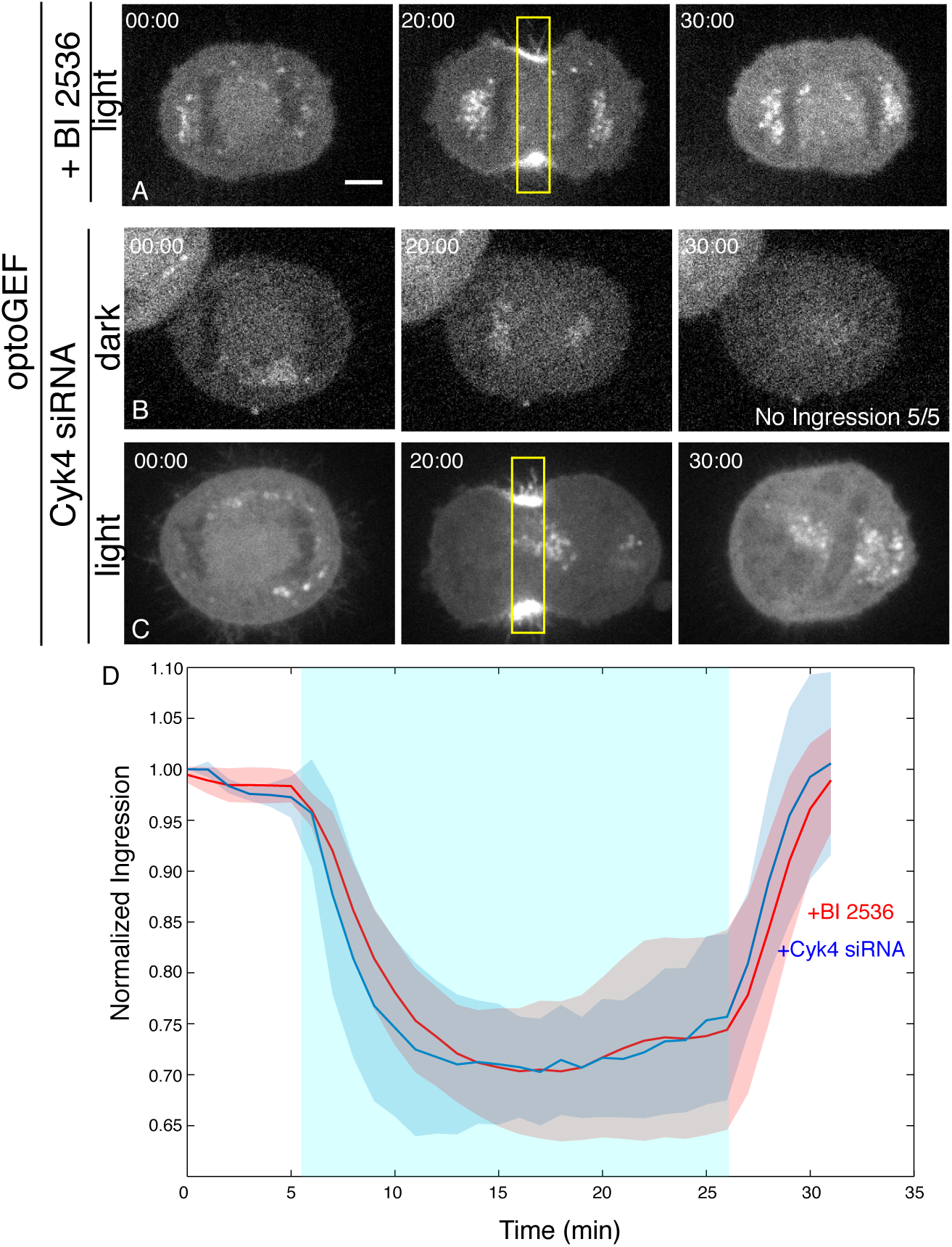
OptoGEF-mediated furrow formation in Cyk4-depleted cells. To generate non-contractile anaphase HeLa cells, cells were either treated with Plk1 inh ibitor BI 2536 (A) or Cyk4 siRNA (B). In the absence of local illumination, BI 2536-treated cells (Fig 2C) and Cyk4 siRNA transfected cells (B) do not form furrows and no ingression at the midzone is observed. Upon Supplemental Figure 2: ROCK inhibition blocks light-mediated furrow formation (A) Following release from nocodazole, HeLa cells were incubated in 10 μM ROCK inhibitor Y27632 for 30 mins. 200 nM BI 2536 was then added to generate non-contractile anaphase cells. Cells were locally illuminated (yellow box) every 20 secs (960 ms pulse) for 20 mins followed by 10 mins without local illumination. Little to no furrow ingression was observed in 4/4 cells. Scale bar is 5 *μ*m. local recruitment of optoGEF (A,C), furrow formation was observed. (D) Average normalized ingression of cells either treated with BI 2536 (red, n=16) or Cyk4 siRNA (blue, n=6). Blue-shaded region designates the photoactivation period during which cells were locally illuminated at the midzone every 20 secs. The mean % Ingression and constriction rate for +BI 2536 treated cells are 34 +/− 13% and 3.7 +/− 1.78 *μ*m/min. The mean % ingression and constriction rate for Cyk4 siRNA transfected cells are 33 +/− 5% and 4.2 +/− 1.76 *μ*m/min. Scale bar is 5 *μ*m.

**Supplemental Figure 4:**
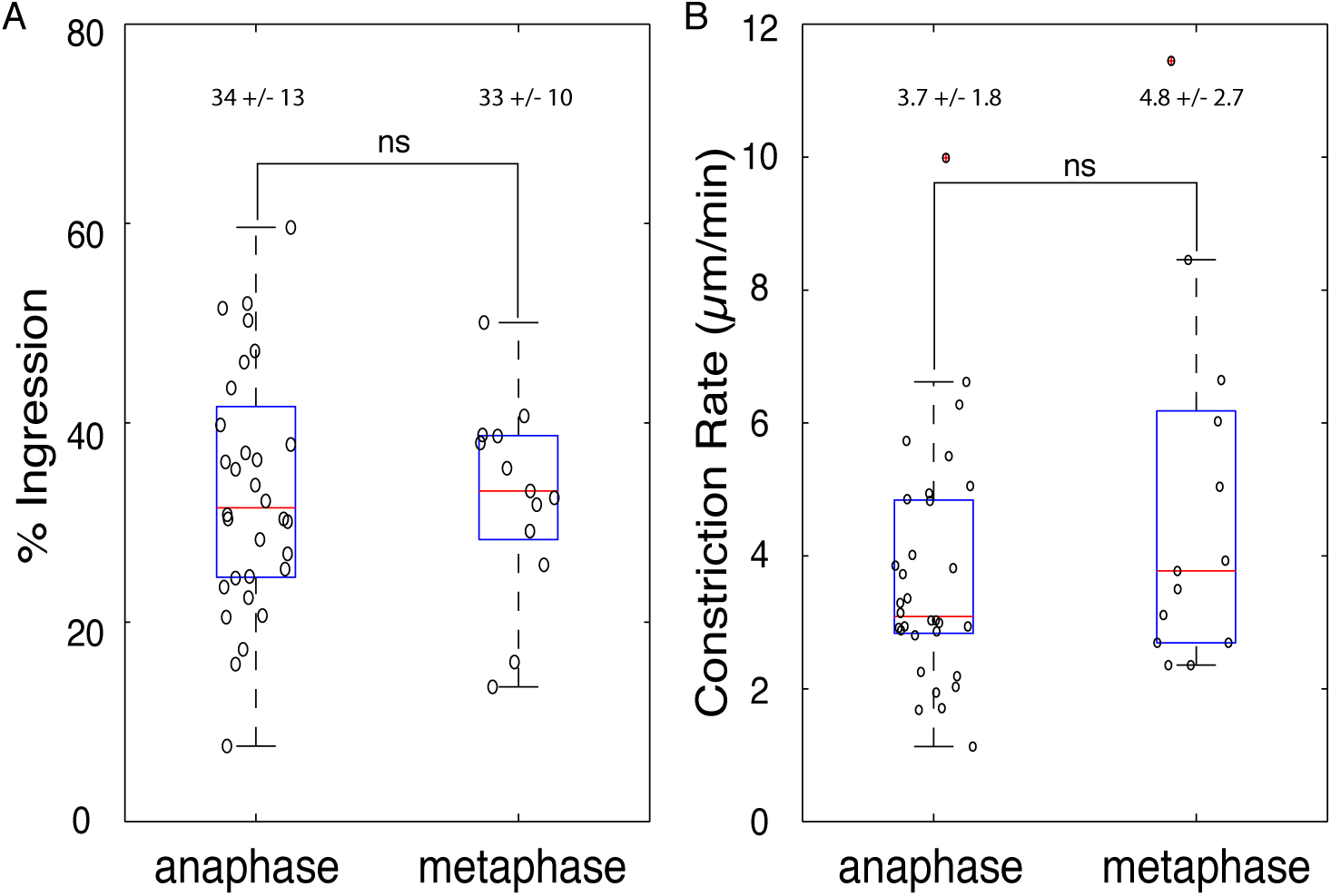
Response to RhoA activation is similar in anaphase and metaphase. Quantification of extent of furrow ingression (A) and rate of constriction (μm/min) (B) of HeLa cells locally illuminated at the midzone during metaphase cells (results with Plk1-inhibited anaphase cells from Figure 3B-C is shown for comparison).

**Supplemental Figure 5:**
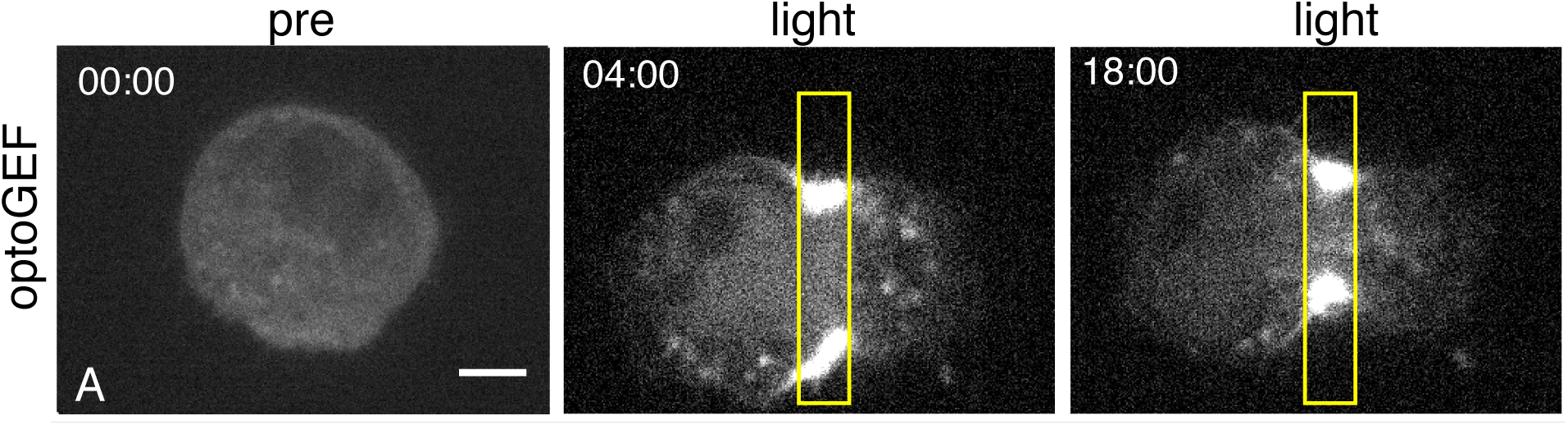
Local activation of RhoA is sufficient to generate furrow formation in nonadherent interphase Hela cells. (A) Images of non-adherent interphase HeLa cells locally illuminated (960ms pulse) every 20 secs for 20 mins. 561 nm images were taken every 20 Supplemental Figure 4: Response to RhoA activation is similar in anaphase and metaphase Quantification of extent of furrow ingression (A) and rate of constriction *μ*m/min) (B) of HeLa cells locally illuminated at the midzone during metaphase cells (results with Plk1-inhibited anaphase cells from Figure 3B-C is shown for comparison). secs. For n=4 cells, the mean % Ingression was 69 +/− 6.3% and the mean constriction rate was 7.34 +/− 4.59 μm/min. The increased adhesive nature of NIH3T3 cells when replated onto glass coverslips stabilized cells for imaging, while HeLa cells frequently moved out of the imaging field. Scale bar is 5 *μ*m.

**Figure S6:**
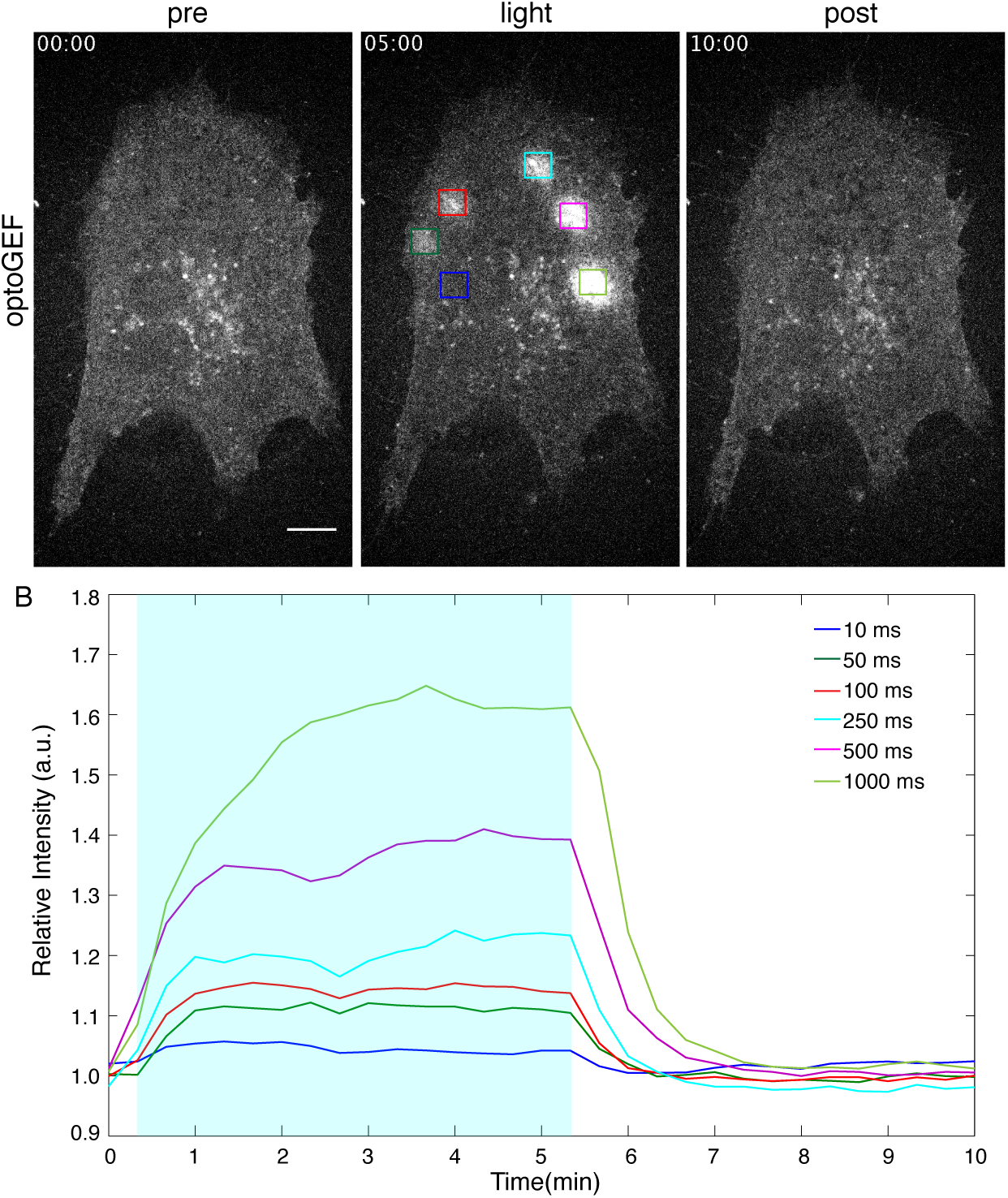
OptoGEF recruitment as a function of the duration of the illumination pulse. (A) Images of an NIH3T3 cell expressing optoGEF just prior to (pre), during (light), and following (post) simultaneous local illumination at the designated regions (boxes) for varying pulse lengths. (B) Quantification of relative intensity of each region as a function of time. Cells were locally illuminated with 405 nm light every 20 secs for designated pulse length for the designated photoactivation period (blue-shaded region) and 561 nm images were taken every 20 secs. Scale bar is 10 *μ*m.

## Supplemental Videos

**Please access the videos here: http://glotzerlab.uchicago.edu/wagnervideos.html**

**Video 1: Light-mediated membrane recruitment of optoGEF.**

A NIH3T3 cell expressing Stargazin-GFP-LOVpep and optoGEF was locally photoactivated with 405 nm light every 20 sec with a 960 ms pulse (yellow box). To visualize optoGEF recruitment, 561 nm images were taken every 20 sec before (5 min), during (5 min), and post (5min) photoactivation.

**Video 2: Light-mediated activation of RhoA induces actin polymerization.**

A NIH3T3 cell expressing Stargazin-GFP-LOVpep, optoGEF^YFP^, and mApple-actin was locally photoactivated with 405 nm light every 20 sec with a 960 ms pulse (yellow box). To visualize the effects of RhoA activation on the actin network, 561 nm images were taken every 20 sec before (15 min), during (15 min), and post (15min) photoactivation.

**Video 3: Light-mediated activation of RhoA induces myosin II accumulation.**

A NIH3T3 cell expressing Stargazin-GFP-LOVpep, optoGEF^YFP^, and mApple-MLC was locally photoactivated with 405 nm light every 20 sec with a 960 ms pulse (yellow box). To visualize the effects of RhoA activation on myosin localization, 561 nm images were taken every 20 sec before (15 min), during (15 min), and post (15min) photoactivation.

**Video 4: Light-mediated activation of RhoA in non-contractile anaphase HeLa cell induces furrow formation at the midzone.**

A non-contractile anaphase HeLa cell (200 nM BI 2536) expressing Stargazin-GFP-LOVpep and optoGEF was locally photoactivated with 405 nm light every 20 sec with a 960 ms pulse (yellow box) at the midzone. To visualize optoGEF recruitment and furrow induction, 561 nm images were taken every 1 min during (20 min) and post (6 min) photoactivation.

**Video 5: Light-mediated activation of RhoA in non-contractile anaphase HeLa cell induces furrow formation at the poles.**

A non-contractile anaphase HeLa cell (200 nM BI 2536) expressing Stargazin-GFP-LOVpep and optoGEF was locally photoactivated with 405 nm light every 20 sec with a 960 ms pulse (yellow box) at the poles. To visualize optoGEF recruitment and furrow induction, 561 nm images were taken every 20 sec during (23 min) and post (14 min) photoactivation.

**Video 6: Simultaneous photoactivation of midzone and polar regions induces a similar response in non-contractile anaphase HeLa cells.**

A non-contractile anaphase HeLa cell (200 nM BI 2536) expressing Stargazin-GFP-LOVpep and optoGEF was simultaneously photoactivated with 405 nm light every 20 sec with a 960 ms pulse (yellow box) at both the midzone and polar regions. To visualize optoGEF recruitment and furrow induction, 561 nm images were taken every 20 sec during (20 min) and post (5 min) photoactivation.

**Video 7: Exogenous activation of RhoA in the poles competes with endogenous furrow formation.**

A dividing HeLa cell expressing Stargazin-GFP-LOVpep and optoGEF was locally photoactivated with 405 nm light every 20 sec with a 960 ms pulse (yellow box) at the poles. To visualize optoGEF recruitment and the effect on endogenous furrow formation, 561 nm images were taken every 20 sec during (30 min) photoactivation.

**Video 8: Light-mediated activation of RhoA induces furrow formation during metaphase.**

A metaphase HeLa cell expressing Stargazin-GFP-LOVpep and optoGEF was locally photoactivated with 405 nm light every 20 sec with a 960 ms pulse (yellow box). To visualize optoGEF recruitment and furrow induction, 561 nm images were taken every 20 sec during (23 min) and post (14 min) photoactivation.

**Video 9: Light-mediated activation of RhoA induces rapid and nearly complete furrow formation in non-adherent interphase NIH3T3 cells.**

A non-adherent interphase NIH3T3 cell expressing Stargazin-GFP-LOVpep and optoGEF was locally photoactivated with 405 nm light every 20 sec with a 960 ms pulse (yellow box). To visualize optoGEF recruitment and furrow induction, 561 nm images were taken every 1 min during (20 min) and post (10 min) photoactivation.

**Video 10: Low levels of global RhoA activation dampen the extent and rate of furrow induction upon local RhoA activation in non-adherent interphase NIH3T3 cells.**

A non-adherent interphase NIH3T3 cell expressing Stargazin-GFP-LOVpep and optoGEF was photoactivated both globally (10 ms pulse) and locally (960 ms pulse, yellow box) with 405 nm light every 20 sec. To visualize optoGEF recruitment and furrow induction, 561 nm images were taken every 20 sec during (10 min) and post (6 min) photoactivation.

## References

Adams, R.R., M. Carmena, and W.C. Earnshaw. 2001. Chromosomal passengers and the (aurora) ABCs of mitosis. Trends Cell Biol. 11:49–54.

Basant, A., S. Lekomtsev, Y.C. Tse, D. Zhang, K.M. Longhini, M. Petronczki, and M. Glotzer. 2015. Aurora B Kinase Promotes Cytokinesis by Inducing Centralspindlin Oligomers that Associate with the Plasma Membrane. Dev Cell. 33:204–215. doi:10.1016/j.devcel.2015.03.015.

Bement, W.M., M. Leda, A.M. Moe, A.M. Kita, M.E. Larson, A.E. Golding, C. Pfeuti, K.-C. Su, A.L. Miller, A.B. Goryachev, and G. von Dassow. 2015. Activator-inhibitor coupling between Rho signalling and actin assembly makes the cell cortex an excitable medium. Nat Cell Biol. 17:1471–1483. doi: 10.1038/ncb3251.

Burkard, M.E., J. Maciejowski, V. Rodriguez-Bravo, M. Repka, D.M. Lowery, K.R. Clauser, C. Zhang, K.M. Shokat, S.A. Carr, M.B. Yaffe, and P.V. Jallepalli. 2009. Plk1 self-organization and priming phosphorylation of HsCYK-4 at the spindle midzone regulate the onset of division in human cells. PLoS Biol. 7:e1000111. doi:10.1371/journal.pbio.1000111.

Drechsel, D.N., A.A. Hyman, A. Hall, and M. Glotzer. 1997. A requirement for Rho and Cdc42 during cytokinesis in Xenopus embryos. Curr Biol. 7:12–23.

Fededa, J.P., and D.W. Gerlich. 2012. Molecular control of animal cell cytokinesis. Nat Cell Biol. 14:440–447. doi:10.1038/ncb2482.

Golsteyn, R.M., K.E. Mundt, A.M. Fry, and E.A. Nigg. 1995. Cell cycle regulation of the activity and subcellular localization of Plk1, a human protein kinase implicated in mitotic spindle function. J Cell Biol. 129:1617–1628.

Jaiswal, M., L. Gremer, R. Dvorsky, L.C. Haeusler, I.C. Cirstea, K. Uhlenbrock, and M.R. Ahmadian. 2011. Mechanistic insights into specificity, activity, and regulatory elements of the regulator of G-protein signaling (RGS)-containing Rho-specific guanine nucleotide exchange factors (GEFs) p115, PDZ-RhoGEF (PRG), and leukemia-associated RhoGEF (LARG). J Biol Chem. 286:18202–18212. doi:10.1074/jbc.M111.226431.

Kishi, K., T. Sasaki, S. Kuroda, T. Itoh, and Y. Takai. 1993. Regulation of cytoplasmic division of Xenopus embryo by rho p21 and its inhibitory GDP/GTP exchange protein (rho GDI). J Cell Biol. 120:1187–1195.

Kosako, H., T. Yoshida, F. Matsumura, T. Ishizaki, S. Narumiya, and M. Inagaki. 2000. Rho-kinase/ROCK is involved in cytokinesis through the phosphorylation of myosin light chain and not ezrin/radixin/moesin proteins at the cleavage furrow. Oncogene. 19:6059–6064. doi:10.1038/sj.onc. 1203987.

Liu, X., T. Zhou, R. Kuriyama, and R.L. Erikson. 2004. Molecular interactions of Polo-like-kinase 1 with the mitotic kinesin-like protein CHO1/MKLP-1. J Cell Sci. 117:3233–3246. doi:10.1242/jcs.01173.

Loria, A., K.M. Longhini, and M. Glotzer. 2012. The RhoGAP Domain of CYK-4 Has an Essential Role in RhoA Activation. Curr Biol. 22:213–219. doi:10.1016/j.cub.2011.12.019.

Lowery, D.M., K.R. Clauser, M. Hjerrild, D. Lim, J. Alexander, K. Kishi, S.-E. Ong, S. Gammeltoft, S.A. Carr, and M.B. Yaffe. 2007. Proteomic screen defines the Polo-box domain interactome and identifies Rock2 as a Plk1 substrate. EMBO J. 26:2262–2273. doi:10.1038/sj.emboj.7601683.

M. Petronczki, M. Glotzer, N. Kraut, and J.-M. Peters. 2007. Polo-like kinase 1 triggers the initiation of cytokinesis in human cells by promoting recruitment of the RhoGEF Ect2 to the central spindle. Dev Cell. 12:713–725. doi:10.1016/j.devcel.2007.03.013.

Meyer, E.J., A. Ikmi, and M.C. Gibson. 2011. Interkinetic nuclear migration is a broadly conserved feature of cell division in pseudostratified epithelia. Curr Biol. 21:485–491. doi:10.1016/j.cub.2011.02.002.

Neef, R., C. Preisinger, J. Sutcliffe, R. Kopajtich, E.A. Nigg, T.U. Mayer, and F.A. Barr. 2003. Phosphorylation of mitotic kinesin-like protein 2 by polo-like kinase 1 is required for cytokinesis. J Cell Biol. 162:863–875. doi:10.1083/jcb.200306009.

Niiya, F., T. Tatsumoto, K.S. Lee, and T. Miki. 2006. Phosphorylation of the cytokinesis regulator ECT2 at G2/M phase stimulates association of the mitotic kinase Plk1 and accumulation of GTP-bound RhoA. Oncogene. 25:827–837. doi:10.1038/sj.onc.1209124.

Otomo, T., C. Otomo, D.R. Tomchick, M. Machius, and M.K. Rosen. 2005. Structural basis of Rho GTPase-mediated activation of the formin mDia1. Mol Cell. 18:273–281. doi:10.1016/j.molcel.2005.04.002.

Piekny, A.J., and M. Glotzer. 2008. Anillin is a scaffold protein that links RhoA, actin, and myosin during cytokinesis. Curr Biol. 18:30–36. doi:10.1016/j.cub.2007.11.068.

Rappaport, R. 1985. Repeated furrow formation from a single mitotic apparatus in cylindrical sand dollar eggs. J Exp Zool. 234:167–171. doi:10.1002/jez.1402340120.

Skelton, N.J., M.F.T. Koehler, K. Zobel, W.L. Wong, S. Yeh, M.T. Pisabarro, J.P. Yin, L.A. Lasky, and S.S. Sidhu. 2003. Origins of PDZ domain ligand specificity. Structure determination and mutagenesis of the Erbin PDZ domain. J Biol Chem. 278:7645–7654. doi:10.1074/jbc.M209751200.

Stewart, M.P., J. Helenius, Y. Toyoda, S.P. Ramanathan, D.J. Muller, and A.A. Hyman. 2011. Hydrostatic pressure and the actomyosin cortex drive mitotic cell rounding. Nature. 469:226–230. doi:10.1038/nature09642.

Strickland, D., Y. Lin, E. Wagner, C.M. Hope, J. Zayner, C. Antoniou, T.R. Sosnick, E.L. Weiss, and M. Glotzer. 2012. TULIPs: tunable, light-controlled interacting protein tags for cell biology. Nat Methods. 9:379–384. doi:10.1038/nmeth.1904.

Tatsumoto, T., X. Xie, R. Blumenthal, I. Okamoto, and T. Miki. 1999. Human ECT2 is an exchange factor for Rho GTPases, phosphorylated in G2/M phases, and involved in cytokinesis. J Cell Biol. 147:921–928.

Toettcher, J.E., O.D. Weiner, and W.A. Lim. 2013. Using optogenetics to interrogate the dynamic control of signal transmission by the Ras/Erk module. Cell. 155:1422–1434. doi:10.1016/j.cell.2013.11.004.

Watanabe, S., Y. Ando, S. Yasuda, H. Hosoya, N. Watanabe, T. Ishizaki, and S. Narumiya. 2008. mDia2 induces the actin scaffold for the contractile ring and stabilizes its position during cytokinesis in NIH 3T3 cells. Mol Biol Cell. 19:2328–2338. doi:10.1091/mbc.E07-10-1086.

Werner, M., E. Munro, and M. Glotzer. 2007. Astral signals spatially bias cortical myosin recruitment to break symmetry and promote cytokinesis. Curr Biol. 17:1286–1297. doi:10.1016/j.cub.2007.06.070.

Wolfe, B.A., T. Takaki, M. Petronczki, and M. Glotzer. 2009. Polo-like kinase 1 directs assembly of the HsCyk-4 RhoGAP/Ect2 RhoGEF complex to initiate cleavage furrow formation. PLoS Biol. 7:e1000110. doi:10.1371/journal.pbio.1000110.

Yüce, O., A. Piekny, and M. Glotzer. 2005. An ECT2-centralspindlin complex regulates the localization and function of RhoA. J Cell Biol. 170:571–582. doi:10.1083/jcb.200501097.

Zanin, E., A. Desai, I. Poser, Y. Toyoda, C. Andree, C. Moebius, M. Bickle, B. Conradt, A. Piekny, and K. Oegema. 2013. A conserved RhoGAP limits M phase contractility and coordinates with microtubule asters to confine RhoA during cytokinesis. Dev Cell. 26:496–510. doi:10.1016/j.devcel.2013.08.005.

Zhang, D., and M. Glotzer. 2015. The RhoGAP activity of CYK-4/MgcRacGAP functions non-canonically by promoting RhoA activation during cytokinesis. elife. 4:204. doi:10.7554/eLife.08898.

